# High temporal resolution RNA-seq time course data reveals mammalian lncRNA activation mirrors neighbouring protein-coding genes

**DOI:** 10.1101/2021.08.25.457323

**Authors:** Walter Muskovic, Eve Slavich, Ben Maslen, Dominik C. Kaczorowski, Joseph Cursons, Edmund Crampin, Maria Kavallaris

## Abstract

**Background:** The advent of next-generation sequencing revealed extensive transcription beyond protein-coding genes, identifying tens of thousands of long non-coding RNAs (lncRNAs). Selected functional examples raised the possibility that lncRNAs, as a class, may maintain broad regulatory roles. Compellingly, lncRNA expression is strongly linked with adjacent protein-coding gene expression, suggesting a potential *cis*-regulatory function. Evidence for these regulatory roles may be obtained through careful examination of the precise timing of lncRNA expression relative to adjacent protein-coding genes.

**Results:** Where causal *cis*-regulatory relationships exist, lncRNA activation is expected to precede changes in adjacent target gene expression. Using an RNA-seq time course of uniquely high temporal resolution, we profiled the expression dynamics of several thousand lncRNAs and protein-coding genes in synchronized, transitioning human cells. Our findings reveal lncRNAs are expressed synchronously with adjacent protein-coding genes. Analysis of lipopolysaccharide-activated mouse dendritic cells revealed the same temporal relationship observed in transitioning human cells.

**Conclusion:** Our findings suggest broad-scale *cis*-regulatory roles for lncRNAs are not common. The strong association between lncRNAs and adjacent genes may instead indicate an origin as transcriptional by-products from active protein-coding gene promoters and enhancers.

## BACKGROUND

Large-scale transcriptomic studies, enabled by improvements in total RNA enrichment and high-throughput RNA profiling technologies, unexpectedly revealed extensive transcription outside the boundaries of known protein-coding genes [1–5]. The class of products of this transcription are now known as long non-coding RNAs (lncRNAs). Throughout the human genome, tens of thousands of these transcripts have been accurately annotated [6, 7]. Despite their ubiquity, the biological significance of most lncRNAs remains unknown. Three consistently documented properties of these transcripts hint at widespread regulatory roles. Firstly, while lncRNA exon sequences are poorly conserved, their promoter region sequences are conserved at levels equivalent to protein-coding genes [3,6,8,9]. Second, lncRNAs display exquisite tissue specificity in their expression patterns [5,6,10]. Thirdly, lncRNA expression is often closely correlated with neighboring protein-coding genes, both in developing [11–13] and adult tissues [6,7,14]. Taken together, these observations indicate lncRNA transcription may promote activation of adjacent, tissue-specific protein-coding genes. Proposed mechanisms to support such broad-scale *cis*-regulatory roles are diverse [15–18].

To test the hypothesis that lncRNAs are ubiquitous *cis*-regulators of gene expression we sought to accurately measure the timing of transcription, a relatively under-studied dimension of regulatory RNA activity. LncRNAs by their nature must be transcribed prior to any *cis*-regulatory role. As transcription is slow relative to the rapid activation of inducible transcription factors, changes in lncRNA expression are expected to precede changes in target gene expression. Indeed, the current limited investigations of lncRNA dynamics in transitioning mammalian cells indicate lncRNA production precedes activation of protein-coding genes [19–21].

Here, we capture with unprecedented temporal resolution the dynamics of several thousand lncRNAs and protein-coding genes in transitioning human cells. Using these data, we demonstrate how differences in transcript production and stability have obscured the sequence of lncRNA and protein-coding gene activation. By accounting for these effects, the high temporal resolution of these data reveal the temporal hierarchy of lncRNA and protein-coding gene activation. Examination of the sequence of events provides insight into the feasibility of broad-scale *cis*-regulatory roles for lncRNAs.

## RESULTS

### Capturing a dynamic transcriptome at high temporal resolution

To capture lncRNA and protein-coding gene transcription dynamics at high temporal resolution, a reliable method to obtain a homogeneous, synchronized cell population was required. To achieve this, we took advantage of the unique growth characteristics of the immortalized human glioblastoma cell line T98G. T98G cells retain growth arrest mechanisms characteristic of untransformed cells [22]. In response to growth factor deprivation, T98G cells undergo reversible G_0_/G_1_ cell cycle arrest. Serum stimulation is sufficient to induce exit from growth arrest, producing a population of tightly synchronized cycling cells, without the need for drug treatment [23–25]. Following stimulation, the transition from quiescence to active cell division is characterized by the induction of a complex transcriptional cascade involving protein synthesis-independent induction of immediate early genes, followed by synthesis-dependent secondary response genes [23]. To capture this transcriptional program at high temporal resolution, synchronized transitioning T98G cells were sampled at 10-minute intervals, from 0 minutes (unstimulated) to 400 minutes (Fig. 1a).

**Figure 1.**
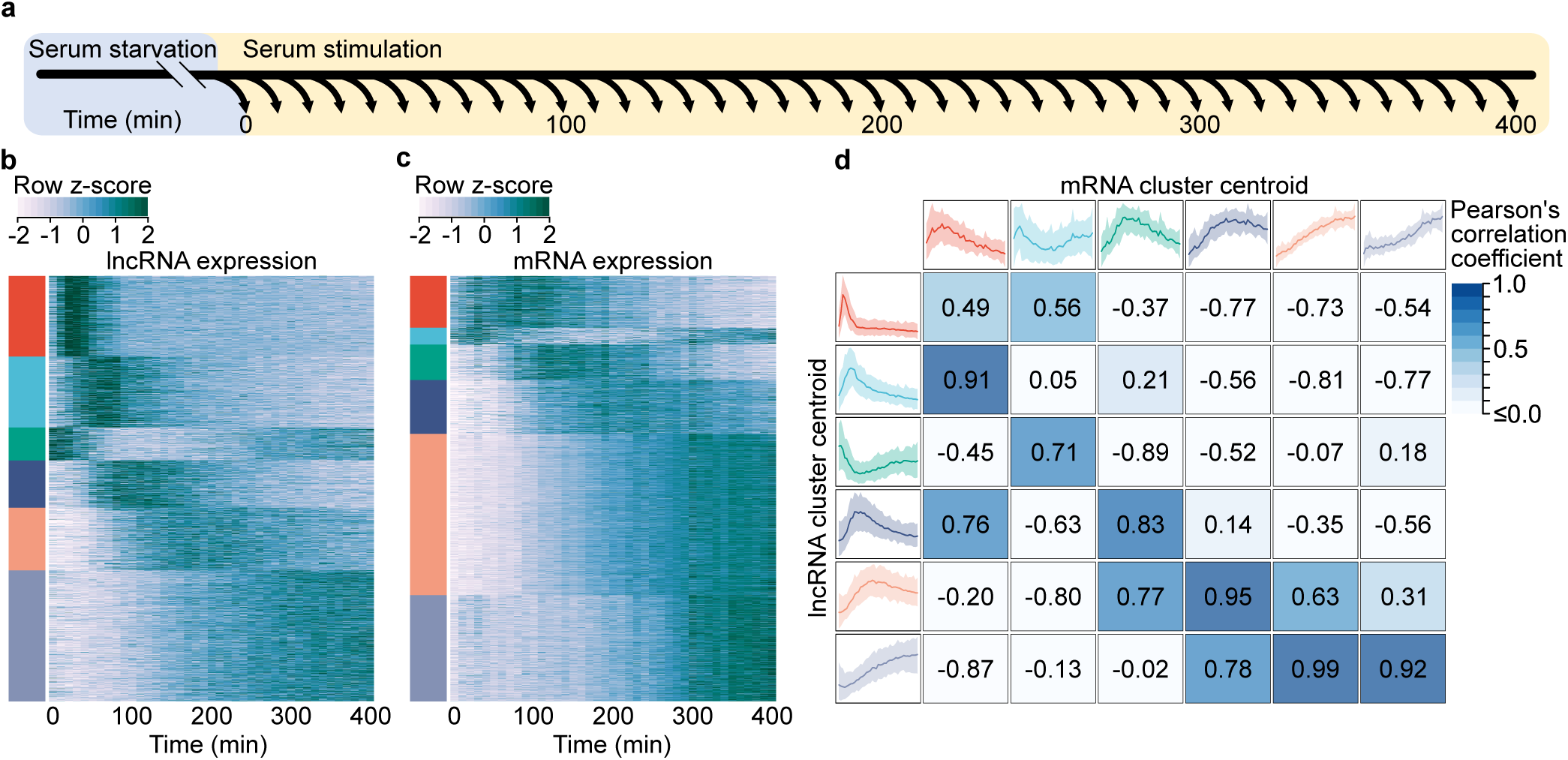
Protein-coding genes and lncRNAs exhibit distinct expression dynamics. **a**, Schematic representation of the experimental design. Following stimulation, cells were harvested at evenly spaced 10-minute intervals, yielding a total of 41 time points. **b**, Heatmap of lncRNA expression. Each row represents an individual z-score normalized lncRNA expression profile. Colored bars indicate six clusters obtained through K-means cluster analysis. **c**, Heatmap of mRNA expression, as in **b**. **d**, Comparison of lncRNA and mRNA cluster centroids. Outer boxes display cluster centroids, capturing the mean expression of all cluster members. Shaded regions representing the 5^th^-95^th^ percentiles of all cluster member expression profiles. Pearson’s correlation coefficients, displayed in the center boxes, were calculated between all lncRNA and mRNA centroid expression profiles.

To obtain gene expression estimates, rRNA-depleted total RNA-seq was performed for all time points. Examination of genome-aligned sequencing reads revealed a large number of lncRNAs were missing from existing genome annotations. To overcome this, *de novo* transcriptome assembly was performed (see Methods), identifying 2803 lncRNAs in addition to 3552 protein-coding genes activated in response to serum stimulation. Of the identified lncRNAs, 33.2% had no overlap with either GENCODE [6] or FANTOM CAT [7] annotated lncRNA transcripts. Notably, many lncRNAs exhibited a rapid increase in expression, peaking within the first 100 minutes of stimulation, followed by an equally rapid decrease in expression (Fig. 1b). In contrast, protein-coding mRNAs displayed more gradual dynamics, with most mRNAs accumulating progressively throughout the time course (Fig. 1c). To directly compare lncRNA and mRNA expression dynamics, we examined the correlation between the prototypical responses displayed by the two transcript classes (Fig. 1d). Notably, coding genes lacked the early rapid response exhibited by most lncRNAs, consistent with previous observations of lncRNAs preceding the expression of protein-coding genes in transitioning mammalian cells [19–21].

However, we noted activated protein-coding genes were significantly longer than the class of lncRNAs (Supplementary Fig. 1). Longer transcription times could introduce delays in mature mRNA accumulation. Protein-coding mRNA half-lives are also known to vary over a wide range, while lncRNAs are generally rapidly degraded by the RNA exosome [26, 27]. The combination of gene length and mRNA stability may mask the time of transcription initiation of protein coding genes (gene activation), impeding accurate comparison with lncRNA activation dynamics. To determine if these effects were obscuring the true protein-coding gene induction times, we next examined the contributions of these two factors to mRNA expression dynamics.

### Transcript stability shapes mRNA expression dynamics

To gain a quantitative understanding of the effect of transcript stability on measured mRNA dynamics we adapted a mathematical model of the transcriptional response proposed by Zeisel et al [28] (see Methods), in which the rate of change of mRNA concentration is determined by a balance between mRNA degradation and the production of new mRNA from unspliced precursor-mRNA (pre-mRNA). RNA-seq reads originating from intronic regions and captured in total RNA-seq have been demonstrated to serve as a useful proxy for nascent transcription [29, 30] and were used to estimate pre-mRNA concentration. Time-invariant splicing and degradation rates were selected that minimized the deviation between model predictions of mRNA concentration relative to measured levels. This model provided a close fit to observed expression dynamics (Fig. 2a-g), enabling estimation of transcript-specific half-lives (Fig. 2h).

**Figure 2.**
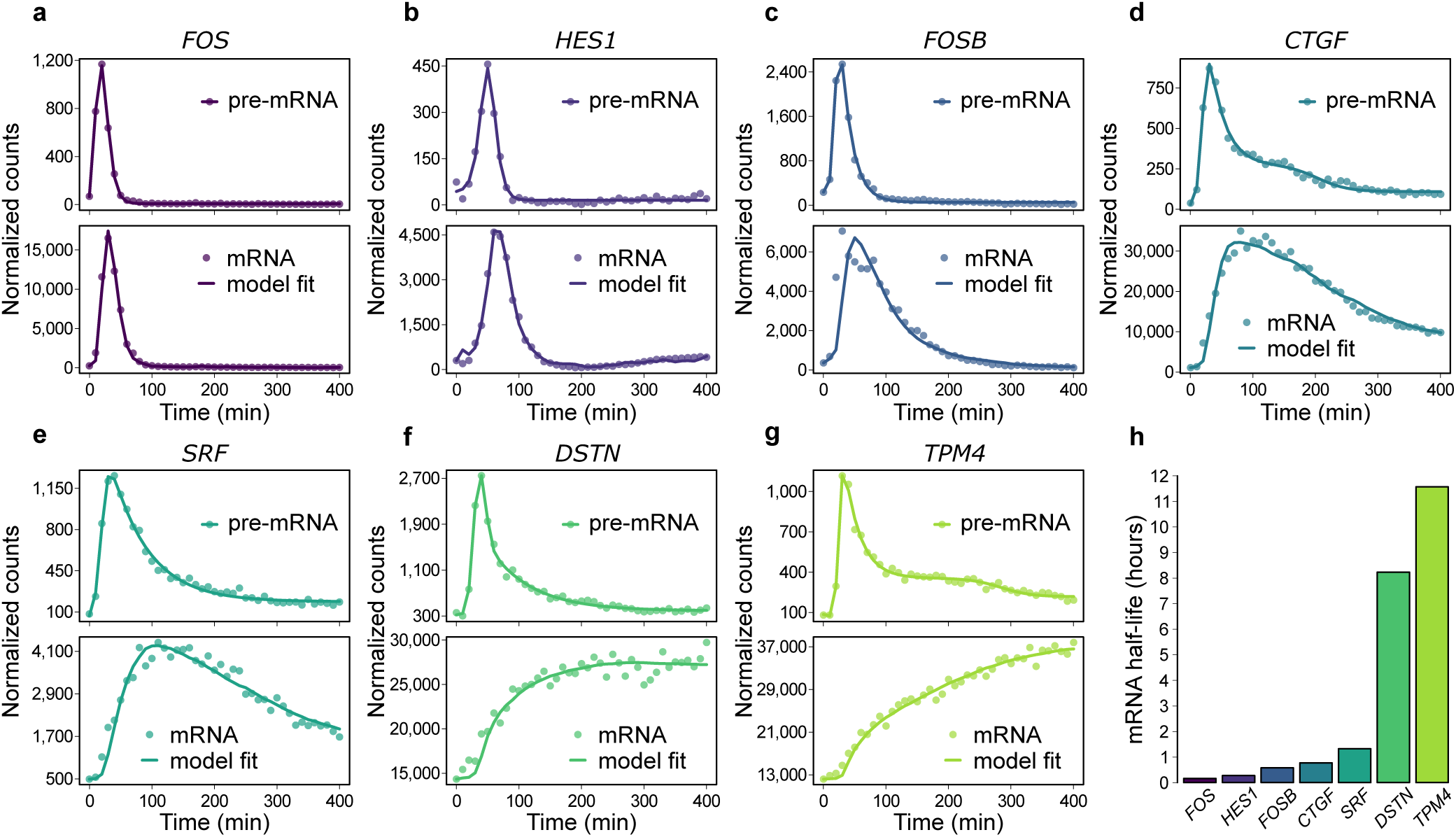
Gene-specific degradation rates shape mRNA dynamics. **a-g**, Pre-mRNA (top panels) and mRNA expression profiles (bottom panels) of seven representative genes with rapid pre-mRNA dynamics. Pre-mRNA and mRNA expression profiles (points) were obtained by quantification of RNA-seq reads mapping to gene introns and exons respectively. Pre-mRNA expression profiles are overlaid with impulse model fits (lines) to aid visualization. mRNA expression profiles are overlaid with the transcription model fits (lines) used to obtain gene-specific mRNA half-lives, presented in **h**.

Genes with relatively unstable mRNA largely recapitulated pre-mRNA dynamics with a short time lag. In contrast, longer mRNA half-lives resulted in expression dynamics increasingly divergent from the transient precursor. These results suggest that for genes encoding stable transcripts, mRNA expression profiles serve as a poor indicator of underlying gene induction dynamics. Furthermore, the confounding effect of transcript stability can be avoided by measuring pre-mRNA expression dynamics for each mRNA transcript through quantification of intron-mapping RNA fragments.

### Gene length introduces RNA production delays

Human gene length varies over a wide range (Supplementary Fig. 1). Protein-coding genes identified in this study ranged from less than a kb to more than a Mb in length, with a mean length of 51.8 kb. In contrast, lncRNAs were observed to be significantly shorter than most protein-coding genes, consistent with previous annotations [6,7,10] with a mean length of 16.6Kb (Supplementary Fig. 1). The time required for Pol II to complete transcript elongation may delay the production of mature mRNA. These effects are expected to be more pronounced for longer genes. This was seen to be the case for the *CACNA1C* gene (Fig. 3a). Visualization of RNA-seq coverage over intronic regions revealed a progressive wave of transcription across the length of the 645 kb gene. Mature mRNA production is correspondingly observed to be delayed by several hours (Fig. 3b). Examination of shorter genes revealed consistent delays in mRNA production due to transcription time (Fig. 3c-e).

**Figure 3.**
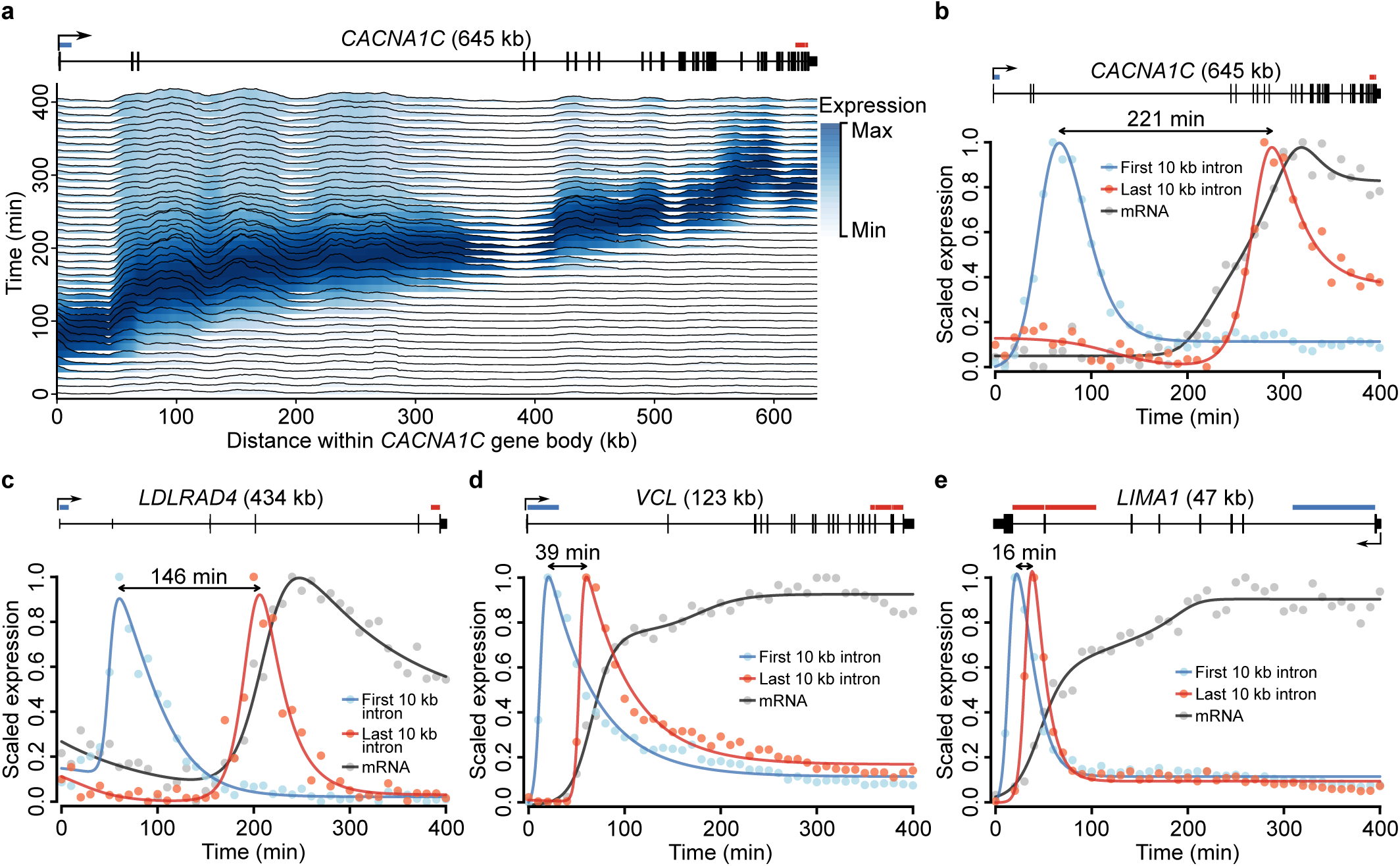
Gene length delays mRNA production. **a**, Transcription across the *CACNA1C* gene body. Ridges display normalized RNA-seq coverage over 1 kb intervals tiled across *CACNA1C* introns. Color intensity indicates the scaled expression of each 1 kb interval across the time course. A right-facing arrow at the 5’ end of the gene schematic (top) indicates the direction of transcription. **b-e**, mRNA and pre-mRNA expression dynamics for four genes of varying length. Pre-mRNA expression is shown for the first and last 10 kb of each gene’s introns, indicated above each gene schematic (top) by blue and red horizontal bars respectively. The approximate delay between transcription of the first and last 10 kb of pre-mRNA is indicated by a left-right arrow between the two expression profile peaks. Expression profiles are overlaid with impulse model fits (lines) and scaled to values between zero and one to facilitate visual comparison.

From these data we estimated transcription elongation to precede at a rate of approximately 2.5 kb/min (Supplementary Fig. 2), in line with previous estimates [31–33]. Assuming this constant rate, the time required to complete transcription elongation of an average length protein-coding gene is approximately 21 minutes. These results suggest mature mRNA expression profiles may be a poor indicator of induction dynamics, particularly for long genes. Further, to negate the effects of transcription delays due to gene length, RNA-seq reads originating from the 5’ end of a gene’s pre-mRNA would be most suitable for determining the timing of gene activation.

### mRNA expression masks underlying gene induction dynamics

Taken together, our findings suggest the combined effects of gene length and transcript-specific degradation rates may combine to mask protein-coding gene induction dynamics. To remove the contributions of these effects, gene expression profiles were quantified for all protein-coding and lncRNA transcripts using only the expression of the first 10 kb of intron sequence. Pre-mRNA profiles (Fig. 4a) revealed protein-coding gene activation is significantly more rapid than indicated by mature mRNA expression levels (Fig. 1c). Within each pre-mRNA expression cluster, genes were ordered by their mRNA expression dynamics (Fig. 4b). Genes with similar pre-mRNA profiles produced a broad range of mature mRNA dynamics, suggesting the combined effects of gene length and transcript stability shape protein-coding gene expression dynamics.

**Figure 4.**
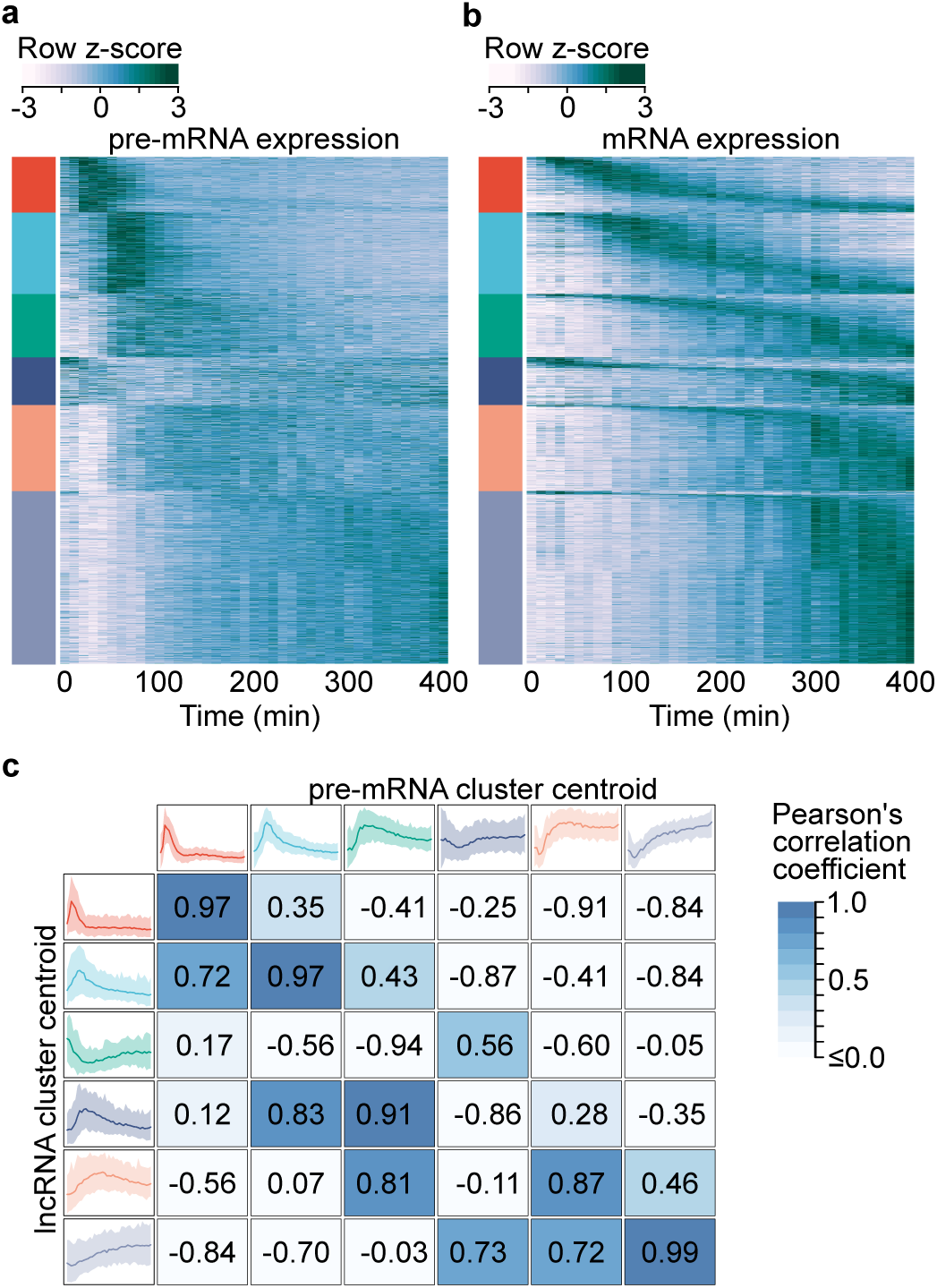
mRNA expression fails to capture gene induction dynamics. **a**, Heatmap of protein-coding gene induction dynamics. Expression profiles were captured using the first 10 kb of gene introns and z-score normalized. Colored bars (left) indicate cluster membership to one of six clusters obtained through K-means cluster analysis. **b**, Heatmap of protein-coding mRNA expression dynamics. Rows within each expression cluster are ranked by the time of peak expression. Rows within **a** and **b** correspond to the same genes. **c**, Comparison of protein-coding gene pre-mRNA and lncRNA expression dynamics. lncRNA cluster centroids (left) are the same as in Fig. 1B, while protein-coding pre-mRNA centroids (top) correspond to the colored bars in **b**. Centroids represent the mean expression of all cluster members, while shaded regions represent the 5^th^-95^th^ percentiles. Pearson correlation coefficients calculated between all lncRNA and protein-coding pre-mRNA centroids are presented.

We next compared the prototypical responses revealed by pre-mRNA with the expression profiles characteristic of lncRNAs (Fig. 4c). In contrast to the relationship implied by mature mRNA expression (Fig. 1d), pre-mRNA dynamics revealed the rapid responses exhibited by lncRNAs are also observed for the induction of protein-coding genes.

### LncRNAs mirror adjacent protein-coding gene expression

Having identified that lncRNAs and protein-coding genes exhibit similar dynamics, we next sought to examine the spatial relationship between lncRNAs and the expression profiles of adjacent protein-coding genes. Before examining the genome-wide relationship, we focused in detail on three well-studied genes activated early in the release from cell cycle arrest (Fig. 5).

**Figure 5.**
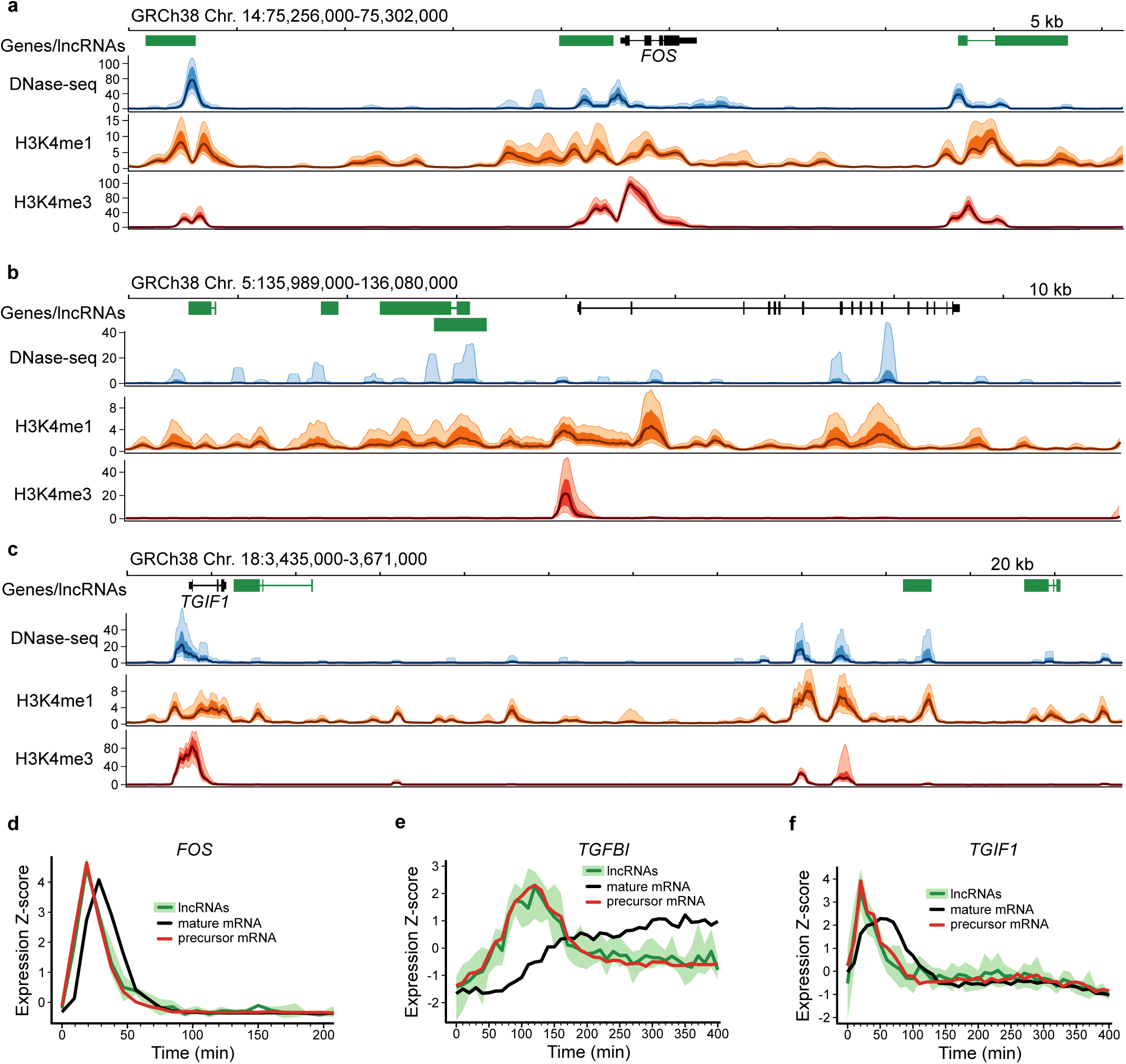
Human lncRNAs mirror adjacent protein-coding gene pre-mRNA expression. **a-c**, NIH Roadmap Epigenomics data for loci surrounding protein coding genes; FOS, TGFBI and TGIF1. A schematic of each loci is presented with GENCODE-annotated protein-coding genes shown in black and lncRNAs in green. NIH Roadmap Epigenomics DNase-seq, H3K4me1 and H3K4me3 histone modification ChIP-seq data from 111 uniformly processed human epigenomes is presented. Lines depict mean -log10(p-value) signal, with dark shaded regions indicating 25%-75% percentiles, and lighter shaded regions the 10%-90% percentiles. **d-f**, Line plots of Z-score normalized protein-coding gene and lncRNA expression values. LncRNA and pre-mRNA was quantified using the expression of the first 10 kb of intronic regions. Mean expression (dark green) and the range of all expression values (shaded light green) is shown for adjacent lncRNAs. Mature mRNA expression is included for comparison.

We first considered the proto-oncogene *FOS*. Following serum stimulation, canonical mitogen-activated protein kinase signaling triggers rapid transcription of immediate early genes, including *FOS* [34]. The encoded transcription factor subunit, c-Fos, dimerizes with c-Jun to form the transcriptional activator AP-1, stimulating further downstream transcriptional changes. Examination of RNA-seq data from the *FOS* locus revealed rapid and transient transcription of *FOS* and two adjacent lncRNAs. Both lncRNAs were associated with regions of increased nuclease sensitivity, revealed by a strong DNase-seq signal across diverse human tissues (Fig. 5a). These regions also overlapped H3K4me1 and H3K4me3 histone marks characteristic of enhancer regions [35, 36]. The expression profiles of both lncRNAs were captured and compared with the adjacent protein-coding *FOS*. Despite the rapid dynamics exhibited within this group, the high temporal resolution of the RNA-seq time series allowed *FOS* pre-mRNA and mRNA dynamics to be separated. Both lncRNAs were found to mirror the expression dynamics of *FOS* pre-mRNA (Fig. 5d).

We next considered *TGFBI*, which encodes an excreted extracellular matrix protein involved in cell adhesion and migration (Fig. 5b). In contrast to the transient dynamics of *FOS*, *TGFBI* exhibited gradual accumulation and increased separation of pre-mRNA and mature mRNA expression profiles (Fig. 5e). Four lncRNAs were identified, clustered upstream of *TGFBI*. Transcription was observed to overlap enhancer-associated chromatin marks. As was observed for *FOS*, comparison of expression dynamics revealed that lncRNA expression mirrored the activation of the adjacent protein coding gene (Fig. 5e).

As a third example, we examined the dynamics of the well-studied transcription factor gene *TGIF1*, which mediates a critical role in attenuating transforming growth factor beta pathway signaling [37]. In addition to the lncRNA antisense to *TGIF1*, two lncRNAs were identified more than 100 kb downstream (Fig. 5c). All lncRNAs overlapped chromatin marks, of variable signal intensity, characteristic of enhancer regions. Consistent with *FOS* and *TGFBI*, analysis of the expression dynamics revealed all lncRNAs mirrored the activation of *TGIF1* (Fig. 5f).

### Protein-coding and lncRNA expression correlation is genome-wide and exhibits synchrony

Close examination of *FOS*, *TGFBI* and *TGIF1* identified adjacent lncRNAs that mirror protein-coding gene activation. To assess the generality of this phenomenon in our data, we next examined the relationship between distance and similarity in expression between all 3552 protein-coding genes and 2803 lncRNAs activated across the human genome. Consistent with observations of individual genes, lncRNAs and protein-coding genes exhibited increasing correlation with increasing genomic proximity (Fig. 6a). As a similar trend is observed within the two transcript classes (Supplementary Fig. 3), a block bootstrap approach was employed (see Methods) to assess uncertainty around the trend between distance and correlation observed between the two transcript classes. Strong deviation of the trend (GAM fit) from the obtained confidence intervals suggests that associations between the expression of lncRNAs and adjacent protein-coding genes is generalizable across our data. To determine whether this trend was consistent between lncRNAs uniquely identified in this study (930) and lncRNAs overlapping existing annotations (1873), the analysis was repeated separately for each group of lncRNAs. The trend between lncRNAs and adjacent protein-coding genes was observed in both groups (Supplementary Fig. 4).

**Figure 6.**
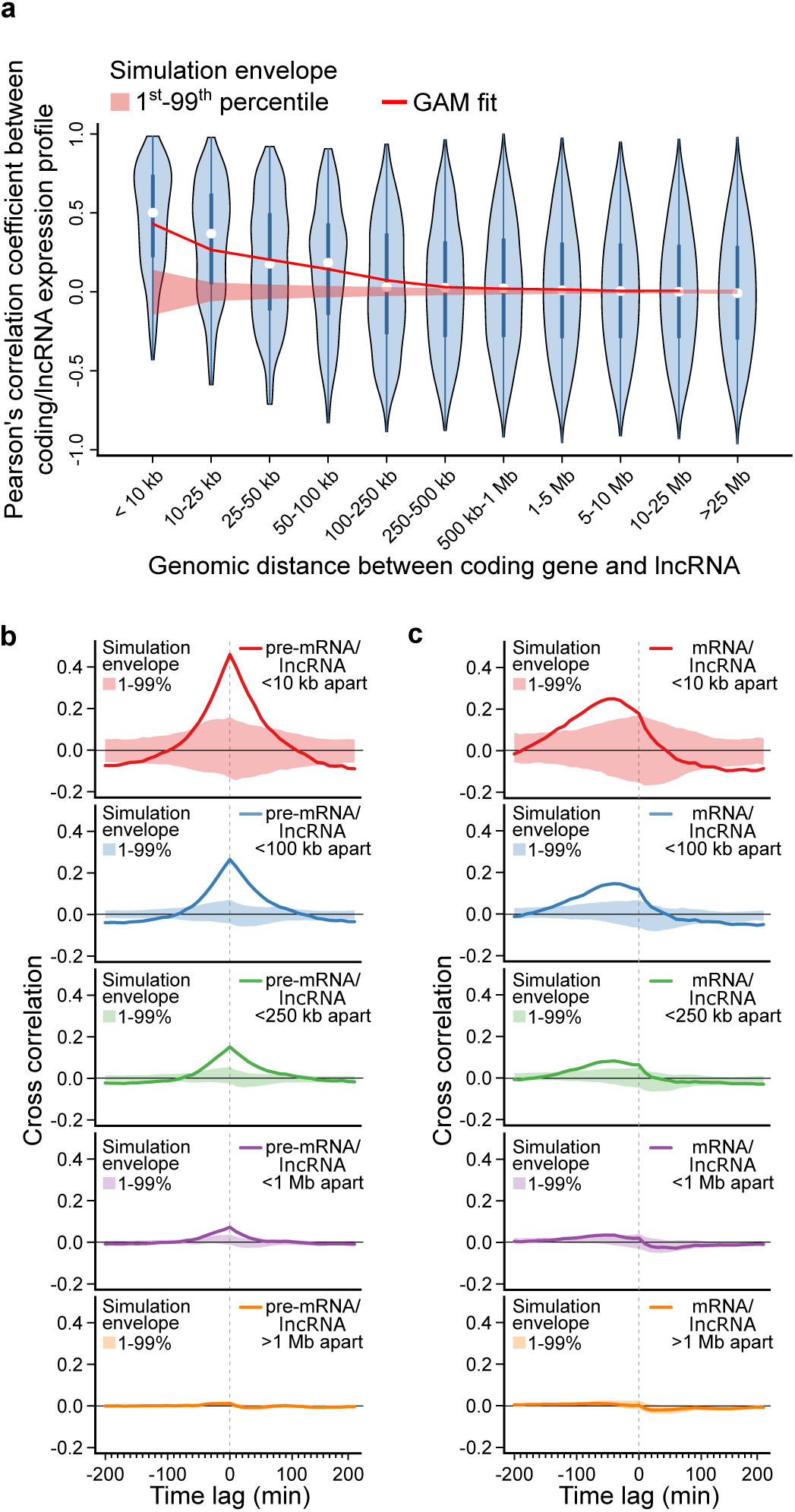
Human lncRNAs mirror adjacent protein-coding gene expression. **a**, Violin plot of Pearson’s correlation coefficients between protein-coding gene and lncRNA expression profiles, binned by distance between transcripts. A generalized additive model (GAM) fit summarizes the relationship between distance and correlation of protein-coding/lncRNA pairs (e.d.f.=8.197, P<2e-16). A simulation envelope, generated using a block-bootstrap approach (see Methods), demonstrates the expected trend under the null hypothesis that distance and correlation are unrelated. The trend in correlation against separation distance lies well outside the simulation envelope indicating a relationship unlikely to be due to chance. Continuous GAM fit and simulation envelope values were overlaid by plotting the mean of each distance bin. **b**, Similarity between expression profiles of coding/lncRNA distance-binned pairs, at time lags of -200 to 200 minutes. Solid lines represent the mean correlation coefficient calculated between distance-binned pairs at varying time-lags of lncRNA expression profiles relative to coding gene expression. Simulation envelopes generated using a block bootstrap approach show the expected cross correlations versus time trends where there is no relationship with separation distance. **c**, produced as in **b**, with coding gene expression profiles replaced with mature mRNA expression, rather than pre-mRNA.

Having identified a genome-wide association between protein-coding gene and adjacent lncRNA expression, we next sought to examine the sequence of events. To determine whether lncRNA expression precedes or trails the activation of adjacent genes, time-lagged lncRNA expression profiles were compared with protein-coding pre-mRNA expression (Fig. 6b). Correlation between lncRNA and protein-coding expression profiles was found to be maximal with a lag of 0 minutes. These results suggest lncRNA expression and coding gene activation are approximately synchronous, consistent with the observations of individual lncRNA-gene pairs (Fig. 5d-f). In contrast, when lncRNA and coding gene dynamics were compared using mature mRNA expression, lncRNA expression appeared to significantly precede protein-coding gene activation (Fig. 6c). These findings highlight the utility of measuring 5’ intron expression to capture gene activation dynamics and provide a possible explanation for the previously reported finding that transcription of lncRNAs precedes protein-coding gene expression [19–21].

### Murine lncRNAs mirror adjacent protein-coding gene expression

In the T98G time series data, simultaneous initiation of lncRNA and adjacent protein-coding expression is consistent across the human genome. To evaluate whether this is also the case in the mouse genome, we examined an RNA-seq time series capturing the immune response of mouse dendritic cells to lipopolysaccharide (LPS) captured at 15 minute time intervals, from 0 to 180 minutes [38]. To identify mouse lncRNAs, *de novo* transcriptome assembly was again performed (see Methods), identifying 1275 lncRNAs and 2882 protein-coding genes activated in response to LPS stimulation. Of the identified lncRNAs, 34.4% had no overlap with GENCODE-annotated lncRNA transcripts.

Consistent with lncRNAs examined in the human T98G time series dataset, mouse lncRNA expression was significantly associated with activation of adjacent protein-coding genes (Fig. 7a). Comparing lagged lncRNA gene expression with nearby protein-coding expression profiles, measured using 5’ intron expression, correlation was again found to be maximal with a time lag of 0 minutes (Fig. 7b). These results suggest synchronous, spatially correlated lncRNA and protein coding gene activation is a general phenomenon in transitioning mammalian cells.

**Figure 7.**
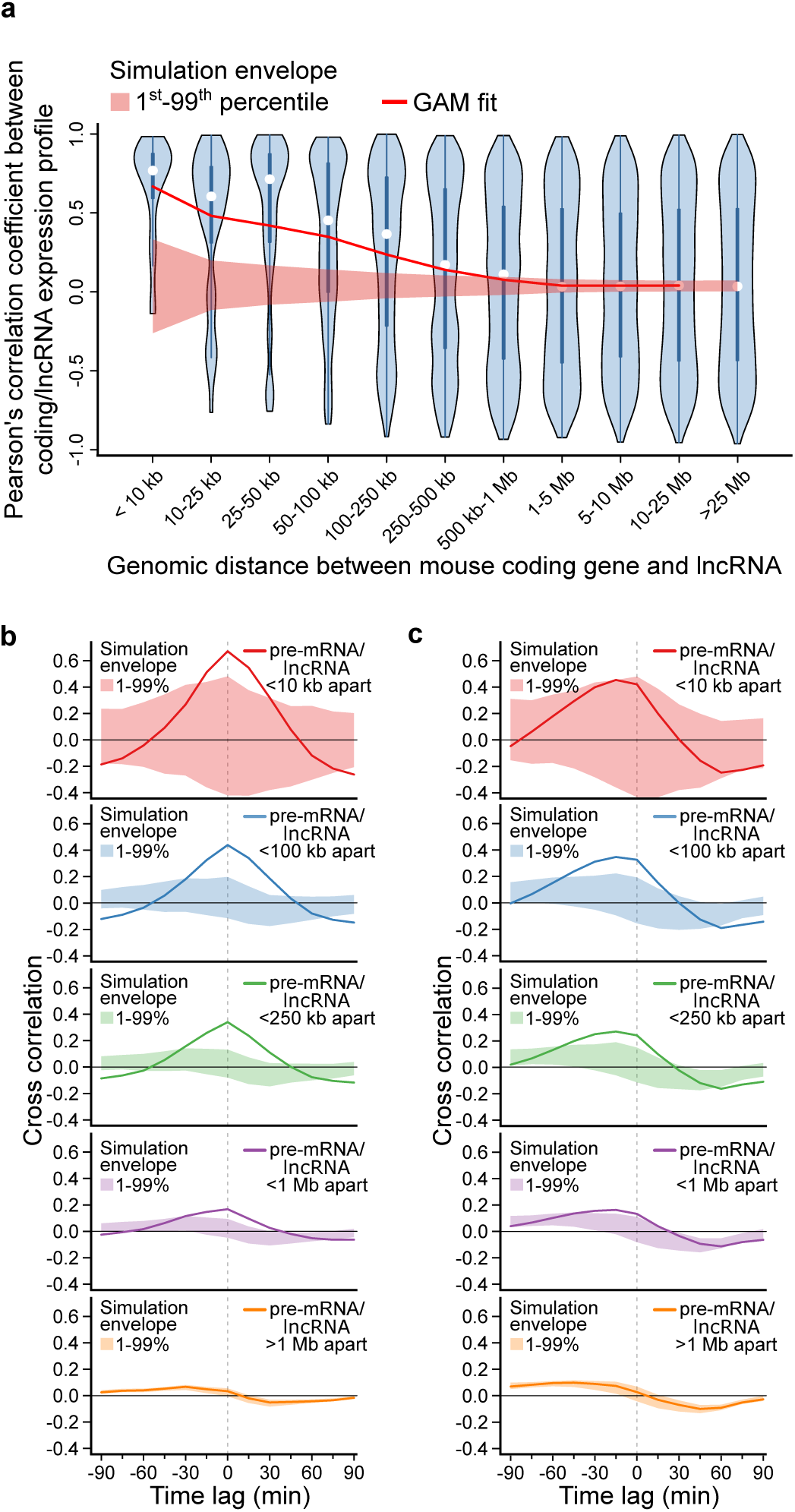
Murine lncRNAs mirror adjacent protein-coding gene expression. Spatial and temporal relationship between protein-coding genes and lncRNAs activated in mouse dendritic cells responding to stimulation with lipopolysaccharide[38].**a**, Violin plot of Pearson’s correlation coefficients between protein-coding gene and lncRNA expression profiles, binned by distance between transcripts. A GAM fit summarizes the relationship between distance and correlation of mouse protein-coding/lncRNA pairs (e.d.f.=7.007, P<2e-16). A simulation envelope, generated using a block-bootstrap approach (see Methods), demonstrates the expected trend under the null hypothesis that distance and correlation are unrelated. The trend in correlation against separation distance lies well outside the simulation envelope indicating a relationship unlikely to be due to chance. Continuous GAM fit and simulation envelope values were overlaid by plotting the mean of each distance bin. **b**, Similarity between expression profiles of coding/lncRNA distance-binned pairs, at time lags of -90 to 90 minutes. Solid lines represent the mean correlation coefficient calculated between distance-binned pairs at varying time-lags of lncRNA expression profiles relative to coding gene expression. Simulation envelopes generated using a block bootstrap approach show the expected cross correlations versus time trends where there is no relationship with separation distance. **c**, produced as in **b**, with coding gene expression profiles replaced with mature mRNA expression, rather than pre-mRNA.

## DISCUSSION

Our findings establish a robust relationship between lncRNAs and the expression of adjacent protein-coding genes. Through genome-wide comparison of lncRNA and coding-gene activation dynamics we have demonstrated that, within the temporal resolution of our measurements, lncRNA and protein-coding gene activation appears to be synchronous.

This observation contrasts with previous reports identifying lncRNA expression to precede activation of protein-coding genes in transitioning mammalian cells [19–21]. Our findings suggest this discrepancy may be attributed to the reliance of previous investigations on measurement of mature mRNA to capture gene expression. We have shown that gene length introduces considerable delays in mRNA accumulation. When combined with differences in transcript stability, our results indicate mRNA levels are an unreliable indicator of gene activation times. In contrast, we have demonstrated that measurement of pre-mRNA expression levels from RNA-seq data reliably captures the timing of gene activation.

Reports of delays between lncRNA and mRNA transcription have been interpreted as evidence supporting functional roles for lncRNAs as pervasive transcriptional regulators [20,21,39]. This reasoning is consistent with non-coding transcripts that must be transcribed prior to any regulatory activity. Where functional regulatory relationships exist, rapid lncRNA expression is expected to occur in advance of changes in target gene expression. Our findings indicate that, with an average length of 16.6 kb and transcription elongation rate of 2.5 kb/min, a typical lncRNA would take 6.6 minutes to be transcribed, excluding the time required for recruitment of regulatory complexes or other proposed functions. The high temporal resolution of the time courses described in this study did not reveal such a delay. Instead, lncRNA and protein-coding gene activation appear to be synchronous.

These findings do not support the existence of broad-scale *cis*-regulatory roles for lncRNAs. Both human and mouse lncRNAs identified in this study arise as transient, low-abundance transcription mirroring adjacent gene activation. These observations are consistent with proposals that the majority of lncRNAs may represent the non-specific initiation of transcription at active regulatory elements [40–42]. Indeed, our findings indicate lncRNAs are associated with chromatin marks characteristic of enhancer elements. This close association of lncRNAs with active enhancers may clarify several observations widely construed as suggestive of biological function. These include the widespread sequence conservation of lncRNA promoter regions [3,6,8,9], strong cell-type and developmental-stage-specific expression [5,6,10] and phenotypic changes observed following ablation of lncRNA loci [43–45]. Sequence conservation of enhancer regions and their regulation of cell-type-specific transcriptional control are well-documented [36, 46]. Conservation of sequence immediately adjacent to lncRNA transcription start sites, previously viewed as lncRNA promoters, may alternatively be interpreted as conserved enhancer regions. Similarly, the characteristic tissue-restricted expression of lncRNAs may reflect activity of the adjacent enhancer. Phenotypes observed following ablation of lncRNA loci may equally be due to loss of underlying regulatory DNA regions, as was recently observed to be the case for a number of zebrafish lncRNAs [47]. Similarly, two recent investigations employing insertion of transcriptional terminator sequences to separate the role of the genomic locus from its RNA products reached similar conclusions [16, 48]. In both cases, *cis* elements were identified as functional, whereas the associated lncRNAs were dispensable.

Importantly, while our observations are consistent with an origin of lncRNAs as transcriptional by-products, they do not preclude potential *trans*-regulatory functions unrelated to activation of adjacent gene expression. These findings also provide an additional criterion by which future studies may distinguish subsets of functional non-coding RNAs. Transcripts that do not originate as transcriptional by-products should be transcribed independent of the activity of neighboring protein-coding gene loci. Further research is required to determine whether independently-regulated lncRNAs are associated with characteristics such as localization with chromatin-associated or gene-silencing factors, increased abundance, stability or sequence-level conservation that may indicate a subset of functional lncRNAs.

## METHODS

### Cell culture and RNA extraction

Human glioblastoma T98G cells obtained from the American Type Culture Collection were cultured in Gibco Dulbecco’s Modified Eagle Medium (DMEM) supplemented with 10% fetal calf serum (FCS) at 37°C in humidified atmosphere with 5% CO2. For each time point two million cells were seeded and allowed to equilibrate for 24 hours, followed by a 72 hour incubation in serum-free DMEM. Cells were stimulated with 20%FCS/DMEM at specified time points, lysed with TRIzol reagent (Ambion), homogenized and frozen for subsequent RNA isolation. RNA extraction and purification was performed using a miRNeasy Mini Kit and RNase-free DNase (Qiagen).

### RNA-sequencing

RNA samples were depleted of ribosomal RNA (rRNA) using Ribo-Zero biotinylated, target-specific oligos (Illumina) combined with RNAClean XP beads (Beckman Coulter). Following purification, rRNA-depleted samples were prepared for sequencing using an Illumina TruSeq Stranded Total RNA library prep kit. After individual library QC, the sample pool size and concentration were determined using a LabChip GX DNA High Sensitivity assay and qPCR using a KAPA Library Quantification Kit (Roche). Uniquely indexed samples were pooled in equimolar concentration, diluted and denatured as one, clustered across eight flow cell lanes and sequenced at 125 bp paired-end resolution using an Illumina HiSeq 2500 v4.0 sequencing system to provide a mean sequencing depth of 37.2 million reads per time point sample.

### Bioinformatic analysis

In addition to the descriptions provided below, all code used to produce the presented analyses and figures, along with links to external data sets are provided in the associated GitHub repository https://github.com/WalterMuskovic/lncRNA_time_course.

### RNA-sequencing data analysis

Sequencing data for the mouse dendritic cell LPS response time course were obtained from NCBI GEO accession GSE56977. A detailed description of the sample preparation and sequencing can be found in the associated publication[38]. Both human glioblastoma T98G and mouse time course reads were trimmed to remove Illumina adapter sequences, with cutadapt, version 1.11 [49]. Trimmed reads were aligned to the GRCh38 and GRCm38 primary genome assemblies using STAR [50], version 2.5.2a. Aligned reads from all timepoints were combined for *de novo* transcriptome assembly with StringTie, version 2.1.3. Read counts were then quantified for each timepoint using the Rsubread R package [51], version 1.34.6. Counts were normalized using the median of ratios method implemented in the DESeq2 R package [52], version 1.24.0. To identify human and mouse genes activated in response to serum stimulation, each gene was tested for autocorrelation using a Ljung– Box test with the stats R package, version 4.0.2. Genes with an adjusted *p*-value cut-off below 0.01 were retained, following correction for multiple-testing with Benjamini-Hochberg adjustment. To assist visualization, protein-coding genes and lncRNAs with similar expression profiles were grouped by K-means cluster analysis. To determine the optimal cluster number (k), the total within-cluster sum of squares (WSS) was calculated for a range of values of k. Examining a curve of WSS according to the number of clusters k, a value was chosen such that adding additional clusters did not greatly reduce the total intra-cluster variation. For all transcript classes a value of k=6 was determined to be appropriate.

### Inference of transcript-specific half-lives

Following the method described by Zeisel et al [28], we model transcription dynamics with the following differential equation:

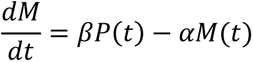

in which the rate change in mRNA concentration 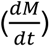 corresponds to the balance between transcription and degradation. *β* denotes the splicing rate coefficient of the pre-mRNA *P*(*t*) to mature mRNA *M*(*t*), which degrades at a rate captured by *α*. Transcript-specific mRNA half-lives are given by 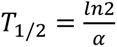. To determine the time-invariant model parameters (*β* and *α*), normalized mRNA and pre-mRNA counts were fit using least squares. To minimize the effects of transcription delays due to gene length, pre-mRNA expression was captured using only reads mapped to the last 10 kb of a gene’s introns. Model parameters were selected as those minimizing the difference between model predictions of mRNA dynamics relative to measured levels.

### Impulse model fits to time course data

To assist with visualization, lines were fit to the pre-mRNA profiles presented in the upper panels of Fig. 2 and the first/last 10 kb of pre-mRNA presented in Fig. 3. Fits were obtained using the parametric impulse model described by Chechik and Koller [53], designed to capture gene expression responses that exhibit an abrupt early response before settling at a second steady-state level. The six-parameter model function described by Chechik and Koller:

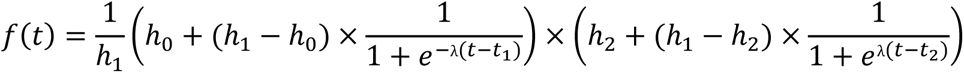

describes two transitions, both with the same slope, captured by *λ*. We generalized the model slightly to allow two transitions with different slopes, defined by *λ*_1_and *λ*_2_:

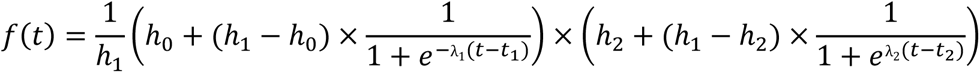

Optimal model parameters were determined by least squares, minimizing the sum of squared error between the impulse model fit and measured pre-mRNA levels.

### Roadmap Epigenomics Project DNase/ChIP-seq data

DNase-seq and histone modification ChIP-seq data for GRCh38 genomic regions were obtained from the NIH Roadmap Epigenomics Project [54]. Data from genomic regions of interest were extracted from genome-wide -log10(p-value) signal tracks containing uniformly processed data from 111 consolidated epigenomes, representing a diverse range of human cell types and tissues [36].

### Block bootstrap

We sought to assess whether coding/lncRNA pairs that are close together are more correlated in their expression profiles than would be expected by chance by plotting a simulation envelope around the relationship between Pearson’s correlation and separation distance to show the 1^st^ and 99^th^ percentiles under the null hypothesis. If the trend is outside the simulation envelope then it indicates there is a relationship between the two that is beyond what is expected by chance. A naive method for the simulation envelope involves creating pseudo samples by randomly permuting the separation distances (but not the Pearson’s correlations) and using these to recreate the “null” trend-where coding/lncRNA correlation and separation distance are not correlated. However, both classes of transcripts are spatially correlated (Supplementary Figure 3) and naive permutation would ignore this dependence. Hence a block bootstrap approach was employed to create the pseudo samples for the simulation envelope [55]. To perform the block bootstrap, pseudo-chromosomes were created by splitting chromosomes into sublengths of a determined block size for each transcript class. Sublengths were then sampled with replacement to obtain the pseudo-chromosomes, with a GAM subsequently fit to the trend in Pearson’s correlation versus separation distance on all the coding/lncRNA pairs in the pseudo-chromosome. A simulation envelope was obtained by taking the 1^st^ and 99^th^ percentiles from 1000 iterations of the block bootstrap. A schematic of the method along with the code used to implement it is provided in the accompanying GitHub repository. To determine the appropriate block size for each transcript class, separation distances were randomly shuffled 1000 times and generalized additive models (GAM) were fit to the relationship between distance and correlation to obtain 1^st^ and 99^th^ quantiles. The distance at which the GAM fit to the unpermuted data exceeded the 99^th^ quantile was taken as the block size, so that the expression profiles between sublengths of chromosome could be considered approximately independent.

### Cross-correlation

The ccf function from the R stats package, version 3.6.1, was used to compute the cross correlation between lncRNA and coding expression profiles, with time lags ranging from -200 to 200 minutes for the T98G time course and -90 to 90 minutes for the mouse LPS time course. The lncRNA expression profile is lagged, while the coding gene expression profile is held constant. To negate any effects of transcription delays due to gene length or transcript half-lives, coding gene pre-mRNA and lncRNA expression was calculated using only the first 10 kb of intron regions. The mean was taken for all coding/lncRNA pairs within the specified separation distance. To gain an estimate of uncertainty in the trend (accounting for autocorrelation in expression profiles along the chromosome), the above procedure was repeated 1000 times on pseudo-chromosomes generated using the block bootstrap method, from which the 1^st^-99^th^ quantiles were obtained in each separation distance category.

## DECLARATIONS

### Ethics approval and consent to participate

Not applicable.

### Consent for publication

Not applicable.

### Availability of data and materials

All data and software used to produce the analyses presented in this work are publicly available. Human glioblastoma T98G RNA-seq time course data have been made available under accession GSE138662. Mouse LPS time course data were obtained from accession GSE56977. All code used to produce the analysis presented in this work are available in the GitHub repository https://github.com/WalterMuskovic/lncRNA_time_course.

### Authors’ contributions

W.M. conceived and planned the project, carried out the in vitro experiments, performed the computational analysis and wrote the manuscript. E.S. devised the methods for analysis of spatial correlation and the block bootstrap. B.M. provided critical input on the analysis of cross-correlation and consulted on the implementation of the block bootstrap method. D.K. performed the RNA-seq library preparation and sequencing. J.C. and E.C. provided input on the bioinformatics analyses and contributed to manuscript revisions. M.K. supervised the project, co-planned experiments, provided critical discussion of the study and contributed to manuscript revisions.

## Acknowledgements

This work was supported by the Children’s Cancer Institute, which is affiliated with the University of New South Wales (UNSW Sydney), and the Sydney Children’s Hospital Network. We would like to acknowledge and thank the donor from whom the glioblastoma T98G cell line used to generate the RNA-seq data described in this publication was derived. We would also like to acknowledge members of the NIH Roadmap Epigenomics Mapping Consortium for generating the described human epigenomics data.

## SUPPLEMENTARY FIGURES

**Supplementary Figure 1.**
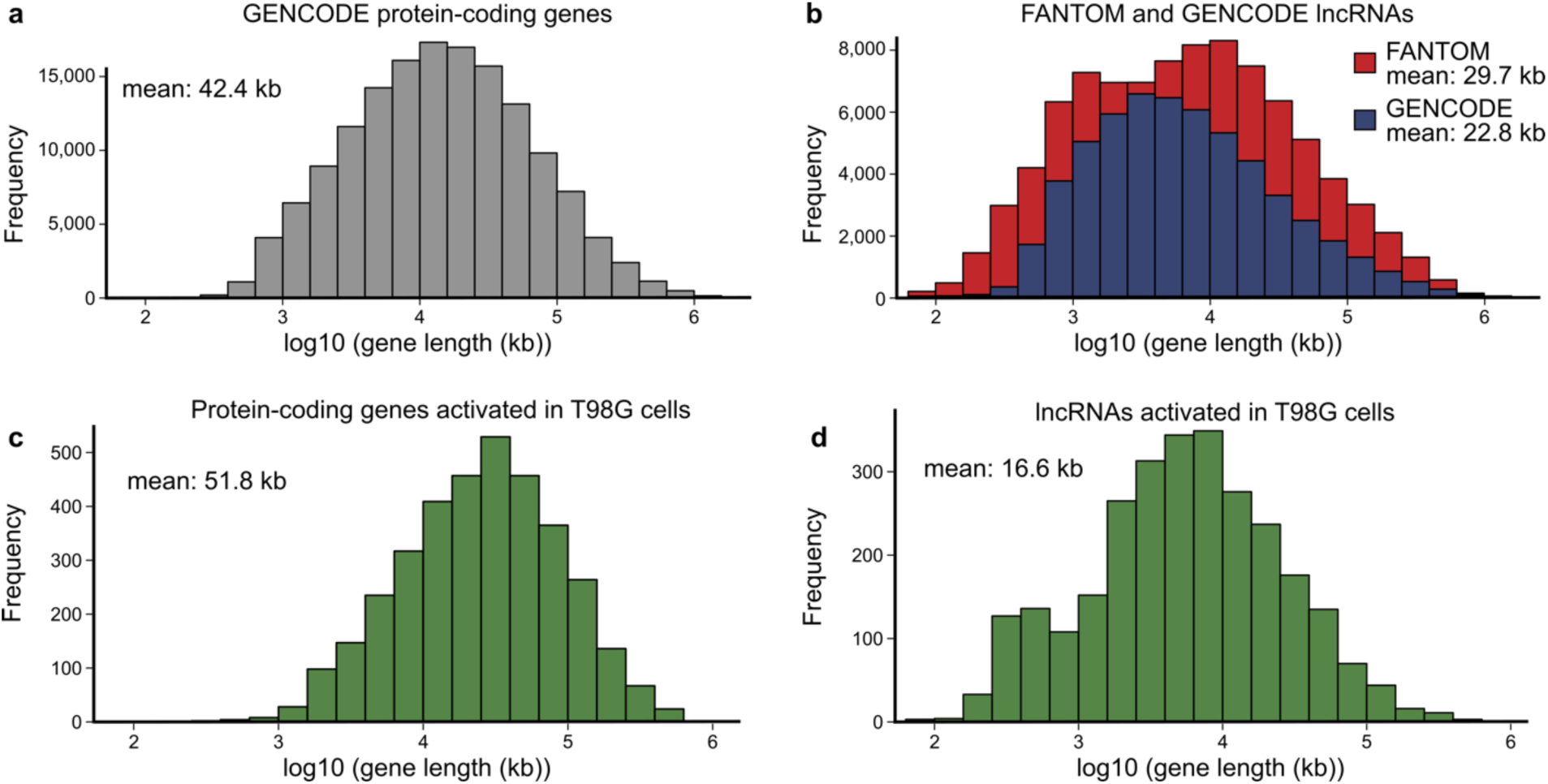
lncRNA and protein-coding gene length. **a**, Histogram showing the distribution of lengths for all protein-coding transcripts in the GENCODE Human Release 29 annotation. **b**, Lengths of all lncRNA transcripts in the FANTOM CAGE associated transcriptome and GENCODE Human Release 29 annotations. **c**, Lengths of all protein-coding genes and **d**, lncRNAs activated in human T98G cells in response to serum stimulation.

**Supplementary Figure 2.**
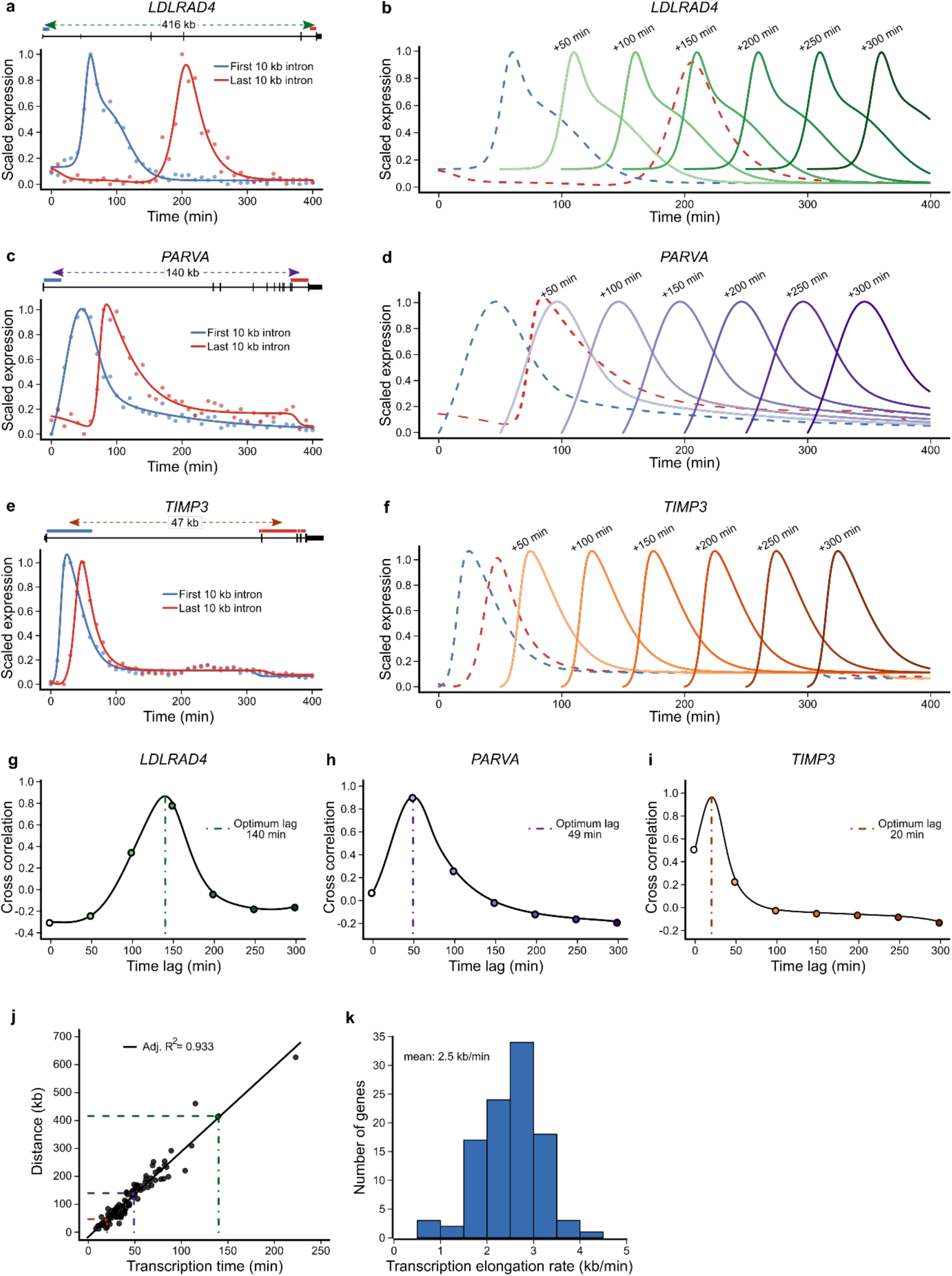
Estimation of the human RNA polymerase II transcription elongation rate. **a**,**c**,**e**, Expression dynamics of the first and last 10 kb of pre-mRNA for three genes of different length. Lines represent impulse model fits to the normalized expression estimates obtained through RNA-seq (points). Included above are schematic illustrations of the three genes *LDLRAD4*, *PARVA* and *TIMP3*, blue and red horizontal bars indicating the regions of intron used to quantify the first and last 10 kb of pre-mRNA respectively. Distance labels indicate the distance between the centers of the first/last 10 kb intervals. **b**,**d**,**f**, Impulse model fits to the pre-mRNA expression dynamics of the three genes as in **a**, **c** and **e** with time-lagged copies of the first 10 kb of pre-mRNA overlaid at intervals of 50 min. **g**-**i**, Lagged correlations between the first and last 10 kb of each gene’s pre-mRNA, obtained by keeping the expression profile of the last 10 kb of pre-mRNA constant and shifting the expression profile of the first 10 kb of pre-mRNA from 0 to 300 min. Filled circles correspond to the time lags overlaid in **b**, **d** and **f**. Vertical lines indicate the time lag at which the correlation between the expression profile of the last 10 kb of pre-mRNA and the lagged expression profile of the first 10 kb of pre-mRNA is maximal. **j**, Scatterplot of the relationship between transcription time and genomic distance with linear model fit overlaid. Colored circles correspond to the transcription times and distances of the three genes presented in **a**-**i**. **k**, Histogram showing the distribution of transcription elongation rates.

**Supplementary Figure 3.**
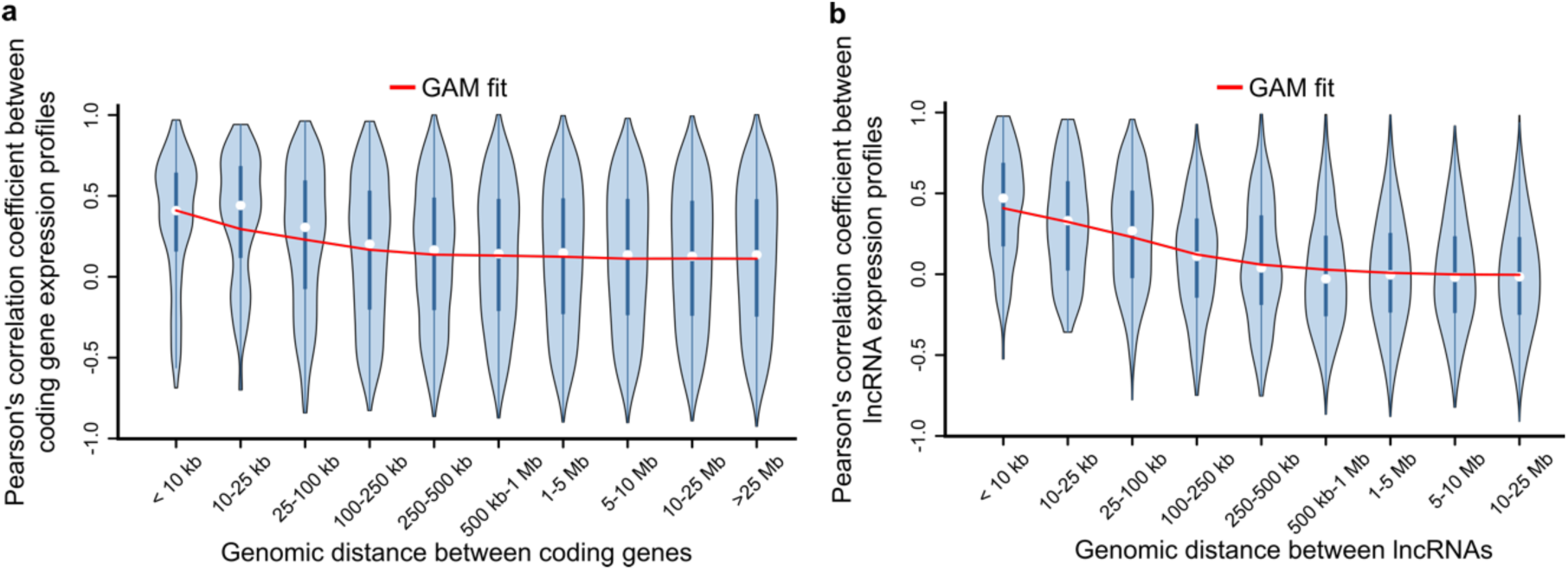
Correlated expression amongst adjacent protein-coding genes and lncRNAs. **a**, Violin plot of Pearson correlation coefficients between protein-coding gene expression profiles, binned by the genomic distance between genes. The overlaid generalized additive model (GAM) fit summarizes the trend between distance and pre-mRNA expression correlation between coding gene pairs (e.d.f=7.703, P<2e-16). **b**, Violin plot of Pearson correlation coefficients between lncRNA expression profiles, binned by the genomic distance between lncRNAs. The overlaid generalized additive model (GAM) fit summarizes the trend between distance and lncRNA expression correlation between lncRNA pairs (e.d.f=8.969, P<2e-16).

**Supplementary Figure 4.**
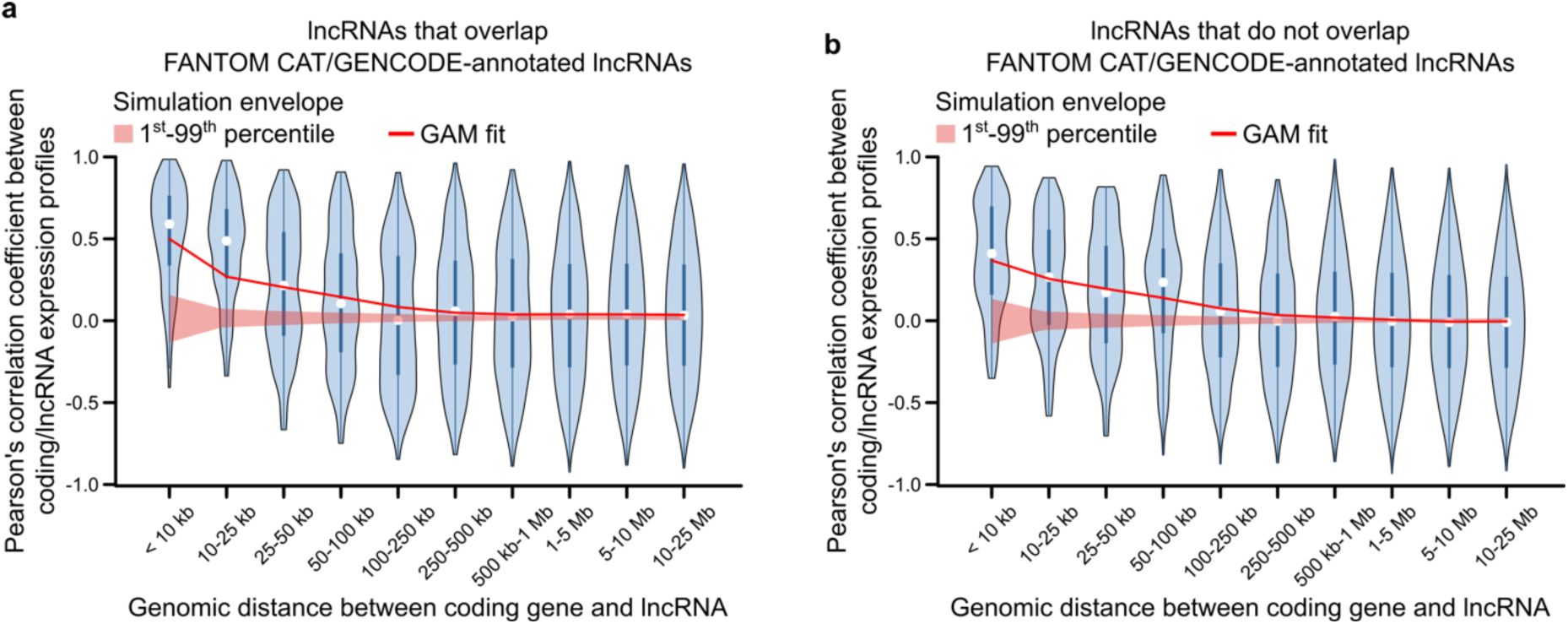
Correlated expression amongst protein-coding genes and lncRNAs with and without overlap of annotated lncRNAs. **a**, Violin plot of Pearson correlation coefficients between the expression profiles of protein-coding genes and lncRNAs that overlap GENCODE/FANTOM CAT- annotated lncRNAs, binned by genomic distance. The overlaid GAM fit summarizes the trend between distance and expression correlation between coding gene/lncRNA pairs (e.d.f=7.851, P<2e-16). **b**, Violin plot of Pearson correlation coefficients between expression profiles of protein-coding gene and lncRNAs that do not overlap annotated lncRNAs. As in **a,** the GAM fit summarizes the trend between distance and expression correlation of the coding gene/lncRNA pairs (e.d.f=7.964, P<2e-16).

